# The ClpX chaperone controls the *Staphylococcus aureus* cell cycle but can be bypassed by β-lactam antibiotics

**DOI:** 10.1101/313668

**Authors:** Kristoffer T. Bæk, Camilla Jensen, Clement Gallay, Niclas Strange Fisker, Ida Thalsø-Madsen, Ana R. Pereira, Wilhelm Paulander, Jan-Willem Veening, Mariana G. Pinho, Dorte Frees

## Abstract

The worldwide spread of *Staphylococcus aureus* strains resistant to almost all β-lactam antibiotics is of major clinical concern. β-lactams interfere with cross-linking of the bacterial cell wall, but the killing mechanism of this important class of antibiotics is not fully understood. Here we show that sub-lethal doses of β-lactams stimulate the growth of *S. aureus* mutants lacking the widely conserved chaperone ClpX. *S. aureus clpX* mutants have a severe growth defect at temperatures below 37°C, and we reasoned that a better understanding of this growth defect could provide novel insights into how β-lactam antibiotics interfere with growth of *S. aureus*. We demonstrate that ClpX is important for coordinating the *S. aureus* cell cycle, and that *S. aureus* cells devoid of ClpX fail to divide, or lyze spontaneously, at high frequency unless β-lactams are added to the growth medium. Super-resolution imaging revealed that *clpX* cells display aberrant septum synthesis, and initiate daughter cell separation prior to septum completion at 30°C, but not at 37°C. FtsZ localization and dynamics were not affected in the absence of ClpX, suggesting that ClpX affects septum formation and autolytic activation downstream of Z-ring formation. Interestingly, β-lactams restored septum synthesis and prevented premature autolytic splitting of *clpX* cells. Strikingly, inhibitors of wall teichoic acid (WTA) biosynthesis that work synergistically with β-lactams to kill MRSA synthesis also rescued growth of the *clpX* mutant, underscoring a functional link between the PBP activity and WTA biosynthesis. The finding that β -lactams can prevent lysis and restore septum synthesis of a mutant with dysregulated cell division lends support to the idea that PBPs function as coordinators of cell division and that β -lactams do not kill *S. aureus* simply by weakening the cell wall.

**Author Summary:** The bacterium *Staphylococcus aureus* is a major cause of human disease, and the rapid spread of *S. aureus* strains that are resistant to almost all β-lactam antibiotics has made treatment increasingly difficult. β-lactams interfere with cross-linking of the bacterial cell wall but the killing mechanism of this important class of antibiotics is still not fully understood. Here we provide novel insight into this topic by examining a defined *S. aureus* mutant that has the unusual property of growing markedly better in the presence of β-lactams. Without β-lactams this mutant dies spontaneously at a high frequency due to premature separation of daughter cells during cell division. Cell death of the mutant can, however, be prevented either by exposure to β-lactam antibiotics or by inhibiting synthesis of wall teichoic acid, a major component of the cell wall in Gram-positive bacteria with a conserved role in activation of autolytic splitting of daughter cells. The finding that the detrimental effect of β-lactam antibiotics can be reversed by a mutation that affect the coordination of cell division emphasizes the idea that β-lactams do not kill *S. aureus* simply by weakening the cell wall but rather by interference with the coordination of cell division.

## Introduction

*Staphylococcus aureus* is a commensal bacterium capable of causing a variety of both localized and invasive infections. Due to its ability to acquire resistance to all relevant antibiotics *S. aureus* remains a major clinical challenge worldwide [1]. The most challenging antimicrobial resistance issue in *S. aureus* has been the dissemination of methicillin-resistant *S. aureus* (MRSA) strains that are resistant to almost all β-lactam antibiotics, one of the safest and most widely used classes of antibiotics ever developed [2]. Early work on the mechanism of action of β-lactams culminated in the discovery that penicillin inhibits crosslinking of peptidoglycan (PG), the central component of bacterial cell walls [3]. The enzymes mediating cross-linking of peptidoglycan strands, the targets of penicillin, were therefore designated penicillin binding proteins (PBPs). The realization that penicillin inhibits PG crosslinking led to the classical model in which penicillin-mediated cell lysis is believed to occur as a consequence of a mechanically weakened cell wall incapable of withstanding high intracellular turgor [3,4]. The killing effect of β-lactam antibiotics, however, has turned out to be more complex [5-9], and may even vary between bacteria, as the organization of PG synthesis and the number of PBPs differ widely between bacterial species [10]. Spherical bacteria such as *S. aureus* have only one cell wall synthesis machine, and *S. aureus* encodes only four PBPs [11]. Notably, MRSA and other Staphylococci have obtained resistance to β-lactams by horizontal acquisition of the *mecA* gene encoding an alternative PBP (PBP2a) that is resistant to inhibition by most β-lactams [12,13]. PBP2a mediated resistance additionally depends on several intrinsic factors that can be targeted by specific compounds to re-sensitize MRSA to β-lactams [14-16]. As an example, inhibitors of wall teichoic acid (WTA) biosynthesis, work synergistically with β-lactams to kill MRSA both *in vitro* and in *in vivo* models of infection, thereby opening a novel paradigm for combination treatment of MRSA [16]. Indeed, a combination strategy pairing β-lactamase inhibitors with β-lactams has proven highly successful in restoring β-lactam efficacy against Gram-negative bacteria [17].

The ClpX chaperone is conserved among bacteria and organelles of eukaryotic cells [18]. The ClpX chaperone has a dual role in cells targeting proteins for degradation by the ClpP protease and, independently of ClpP, facilitating protein folding and interactions [18]. *S. aureus clpX* mutant exhibits a mild growth defect at 37°C that is severely exacerbated at 30°C [19,20]. This cold-sensitive growth defect of the *clpX* mutant is independent of ClpP, and is alleviated by loss-of-function mutations in the *ltaS* gene [20,21]. *ltaS* encodes the LtaS synthetase that is required for synthesis of lipoteichoic acid (LTA), an essential cell wall polymer of Gram-positive bacteria controlling cell division and autolytic activity [22]. Interestingly, inactivation of ClpX restored the septum placement defects of cells depleted for LTA, suggesting a link between ClpX and cell division in *S. aureus* [20].

Indeed, the data presented here demonstrate that ClpX becomes critical for progression of *S. aureus* septum synthesis as the temperature decreases. In cells with delayed septum synthesis, autolytic splitting of daughter cells is activated prior to septum completion resulting in cell lysis, unless β-lactam antibiotics are added to the growth medium. Strikingly, inhibitors of WTA biosynthesis, similarly to mutations in *ltaS* specifically rescue growth of *S. aureus clpX* mutants, emphasizing a fundamental connection between the transpeptidase activity of PBPs and teichoic acids biosynthesis.

In conclusion, this study identifies the ClpX chaperone as an important player in *S. aureus* cell division at sub-optimal temperatures, and provides novel insight into the link between β -lactam antibiotics and cell division in this important pathogen.

## Results

### β-lactam antibiotics stimulate growth of *S. aureus clpX* mutants

Serendipitously, in determining the susceptibility of the *S. aureus clpX* mutant to oxacillin, we repeatedly observed zones of improved growth at a certain distance from the filter discs containing the antibiotic, a phenomenon that was not observed for wild-type strains (marked by arrow in Fig 1a). This observation indicated that sub-lethal concentrations of oxacillin stimulate growth of the *clpX* mutant. Indeed, addition of sub-lethal concentrations of oxacillin rescued the severe growth defect normally seen for *S. aureus clpX* mutants at 30°C (Fig 1b). To investigate if growth of *S. aureus clpX* mutants is generally improved by addition of β-lactam antibiotics, three *S. aureus* strains of clinical origin, representing both MRSA (JE2) and methicillin-sensitive *S. aureus* (SA564 and Newman), and the corresponding *clpX* deletion strains were grown in broth containing oxacillin, meropenem or cefuroxime (representing three different chemical classes of β-lactams) in various concentrations below and above the previously determined MIC values [23]. We found that the presence of β-lactam antibiotics increased the growth rate and the final yield of the *clpX* mutants in all strain backgrounds (Fig 1c and 1d; S1 Fig). As shown previously [23], inactivation of *clpX* increased the MIC values in the JE2 background (S1 fig), but not in the MSSA strain backgrounds. A wide range of β-lactam concentrations was tested, but we did not identify any concentration at which the growth rates of the wild-type strains were enhanced (S1 Fig). For comparison, we also included *clpP* mutants, but observed no or only a minor stimulatory effect on the growth rate of the *clpP* mutants in the presence of β-lactams (S1 Fig). We conclude that the ClpX dependent growth defect that is suppressed by β-lactams is caused by loss of ClpX chaperone activity, not loss of ClpXP protease activity.

**Fig 1.**
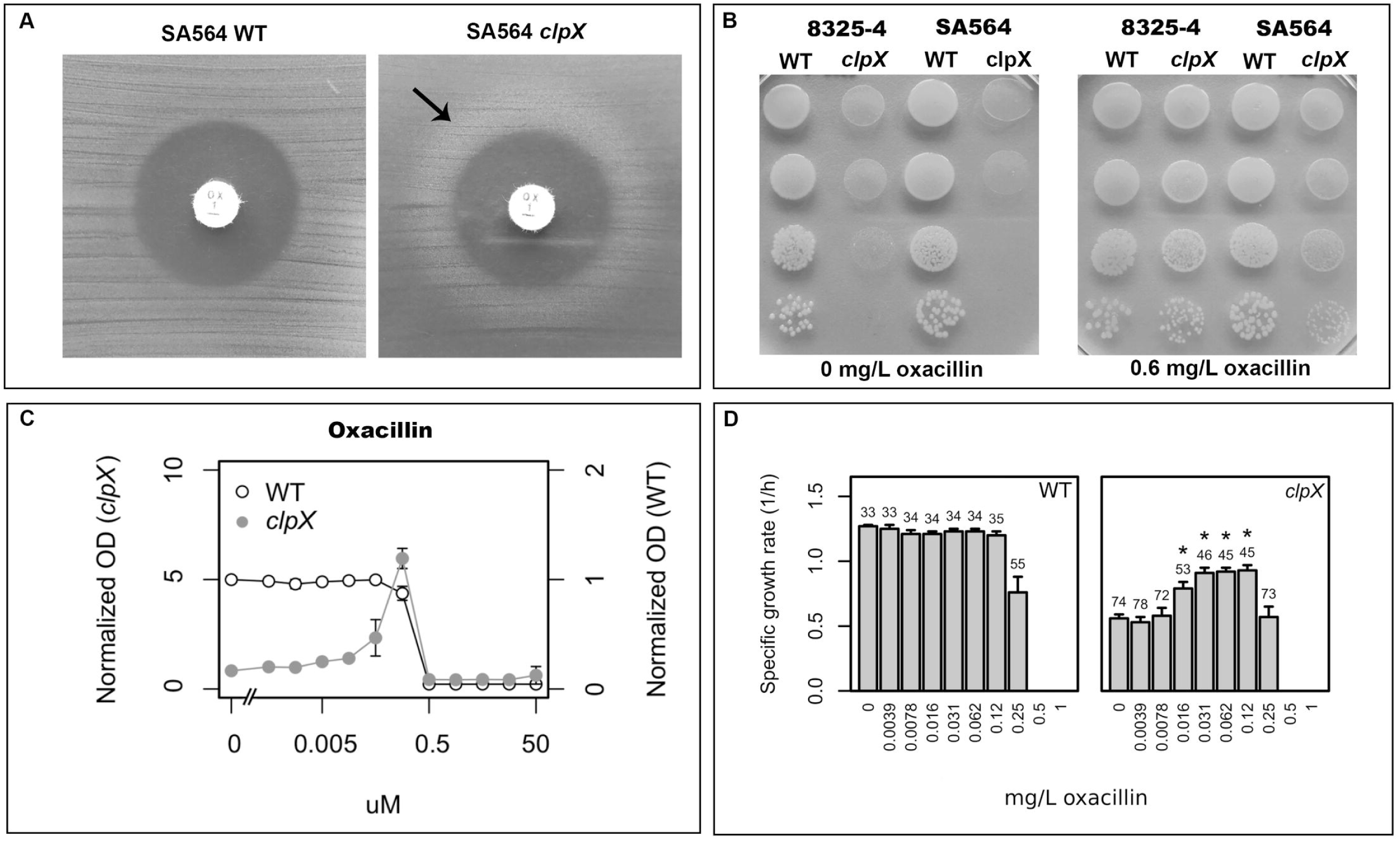
Growth of *S. aureus clpX* mutants is stimulated by β-lactams. (**a**) SA564 wild-type and SA564Δ*clpX* were plated at 37°C and tested for susceptibility to oxacillin in a disc diffusion assay. Disks contain 1 μg oxacillin. (**b**) The *S. aureus* wild-type strains, 8325-4 and SA564 and the corresponding *clpX* deletion mutants were grown exponentially in TSB at 37°C. At OD_600_ = 0.5, the cultures were diluted 10^1^, 10^2^, 10^3^ and 10^4^-fold, and 10 μl of each dilution was spotted on TSA plates +/-oxacillin and the plates were subsequently incubated at 30 °C for 24 h. (**c**) *S. aureus* SA564 wild type and the *clpX* mutant strains were grown overnight at 37°C, diluted 1:200 and grown at 37°C until mid-exponential phase. These cultures were then diluted into TSB containing increasing concentrations of the oxacillin in a 96-well format, and the plates were incubated for 24 h at 30°C. The values represent means of OD values, normalized to the OD values obtained without compound. Error bars indicate standard deviations. Note that different scales were used on the two axes due to the difference in growth between the WT and *clpX* mutant: values for the *clpX* mutant are indicated on the left vertical axis, and values for the WT are indicated on the right vertical axis to allow easy comparison of growth between the two strains. (**d**) Mean growth rates (h^-1^) for *S. aureus* SA564 wild-type and *clpX* when grown at 30°C as described above. Numbers above bars indicate mean doubling time in minutes. The standard error of the mean (error bars) was calculated using values from three biological replicates. Asterisks indicate significantly improved growth rate (P < 0.05). The P values were obtained by comparing the growth rates at each concentration to the growth rate without antibiotics and were calculated using Student’s t-test.

### Oxacillin prevents premature growth arrest and spontaneous lysis of *clpX* cells

Given the unusual ability of β-lactams to stimulate growth of the *clpX* mutant, we reasoned that a better understanding of the *clpX* growth defect could provide novel insights into how β-lactam antibiotics interfere with growth of *S. aureus*. To investigate how β-lactams improve growth of the *clpX* mutant, we studied the growth of single cells of the *S. aureus* SA564 wild type and *clpX* mutant in the absence or presence of oxacillin at 30°C using automated phase contrast time-lapse microscopy. Interestingly, the time-lapse experiments demonstrated that only few *clpX* cells started to divide and that many *clpX* cells lysed spontaneously or stopped dividing early on in the experiment (Fig 2a and S1 Movie); accordingly only about one third of the *clpX* mutant cells ended up forming a micro-colony. While addition of sub-lethal concentrations of oxacillin did not affect growth of the wild type, growth of the *clpX* mutant was clearly stimulated at the single-cell level (Fig 2b). In the presence of oxacillin, only very few *clpX* cells lysed or stopped dividing during the experiment, and the *clpX* mutant reached a higher number of cells than the wild type by the end of the experiment (Fig 2b and 2c). To quantify the cell generation time (time between two divisions), we tracked the fate of each individual cell in one representative micro-colony of each strain during the first 8 h of the experiment (S2 Fig). For the wild type, the average cell generation time was 66 ± 42 min without oxacillin, and 78 ± 58 min with oxacillin, with only few divisions observed after 4 h in both cases (Fig 2c, S2 Fig, and S1 Table). For the *clpX* mutant, the average cell generation time was 71 ± 36 (note, only 11 divisions) min in the absence of oxacillin. Interestingly, in the presence of oxacillin the generation time for *clpX* mutant cells decreased throughout the experiment to an average of 38 ± 23 min during the last 2 h (Fig 2c, S2 Fig, and S1 Table). Hence, oxacillin seems to stimulate growth of the *clpX* mutant by both shortening the generation time and preventing premature growth arrest, and lysis of the mutant.

**Fig 2:**
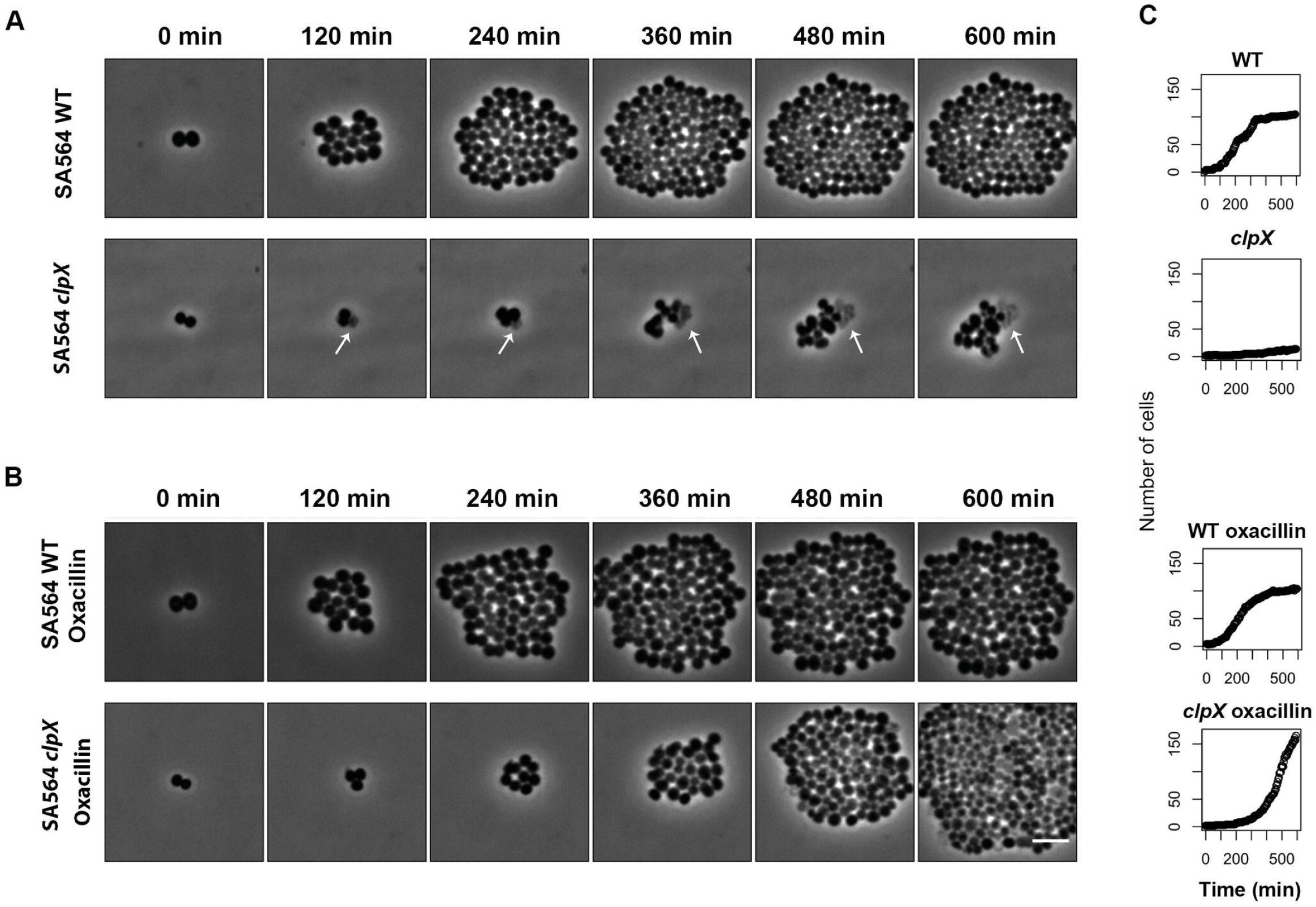
Single cell analysis reveals that oxacillin increases the growth rate and prevents spontaneous lysis of the *S. aureus clpX* mutant. Still images from time-lapse microscopy (phase contrast) of SA564 wild-type and SA564 *clpX* cells growing on a semisolid surface at 30°C, without (**a**) or supplemented with oxacillin at 0.008 μg ml^-1^ (**b**). The still images are taken from movies (see S1 Movie) showing the typical growth of one micro-colony among at least 36 imaged micro colonies (see Methods). Scale bar, 2 μm. (**c**) Number of cells present at each time point in the shown time-lapse image series.

### Aberrant septum synthesis and premature splitting of *clpX* daughter cells

To further investigate the *clpX* phenotypes, we studied the morphology of wild-type and *clpX* mutant cells by transmission electron microscopy (TEM) and scanning electron microscopy (SEM) after growth at 30°C. As reported before, cells lacking ClpX are significantly smaller than wild-type cells and have a thickened cell wall (Fig 3a) [23]. Consistent with the spontaneous cell lysis observed in the time-lapse microscopy approximately 10% of the *clpX* mutant cells grown at 30°C appeared as lysed ghost cells in the TEM images (S3 Fig). Interestingly, these ghost cells had a characteristic appearance in which the cell wall was ripped apart at the tips of ingrowing, still incomplete, septa (see examples in Fig 3c), indicating that these cells underwent lysis while in the process of daughter-cell splitting. To divide, *S*. *aureus* builds a septal cross wall generating two hemispherical daughter cells connected through a narrow peripheral ring [24,25]. Resolution of this peripheral wall ring leads to rapid splitting of daughter cells, in a process designated as “popping” [25]. Popping normally occurs only in cells with closed septa and consistent with this notion, daughter cell splitting was not observed in wild-type cells with incomplete septa (S3 Fig). In contrast, a fraction (quantified below) of the *clpX* mutant cells seemed to have initiated splitting of daughter cells despite having incomplete septa (see examples Fig 3d and 3e, and S3 Fig). In TEM images, initiation of splitting appeared as small invaginations at the external edge of the septum in cells with incomplete septa, but occasionally, a complete splitting of the ingrowing septum and elongation of cells were observed (Fig 3d and 3e, middle panels). In SEM images, the stage of septum ingrowth could not be monitored. Strikingly, however, it was possible to visualize *clpX* cells in the process of splitting despite displaying a non-closed septal cross-wall (seen as a hole; Fig 3c right panel), or *clpX* mutant cells in the process of splitting while still being connected by an undivided cytoplasm (Fig 3d right panel). Taken together, these findings strongly suggest that in the absence of ClpX, the system controlling the onset of autolytic separation of daughter cells becomes dysregulated, and that premature splitting of *clpX* cells with incomplete septa result in cell lysis.

**Fig 3.**
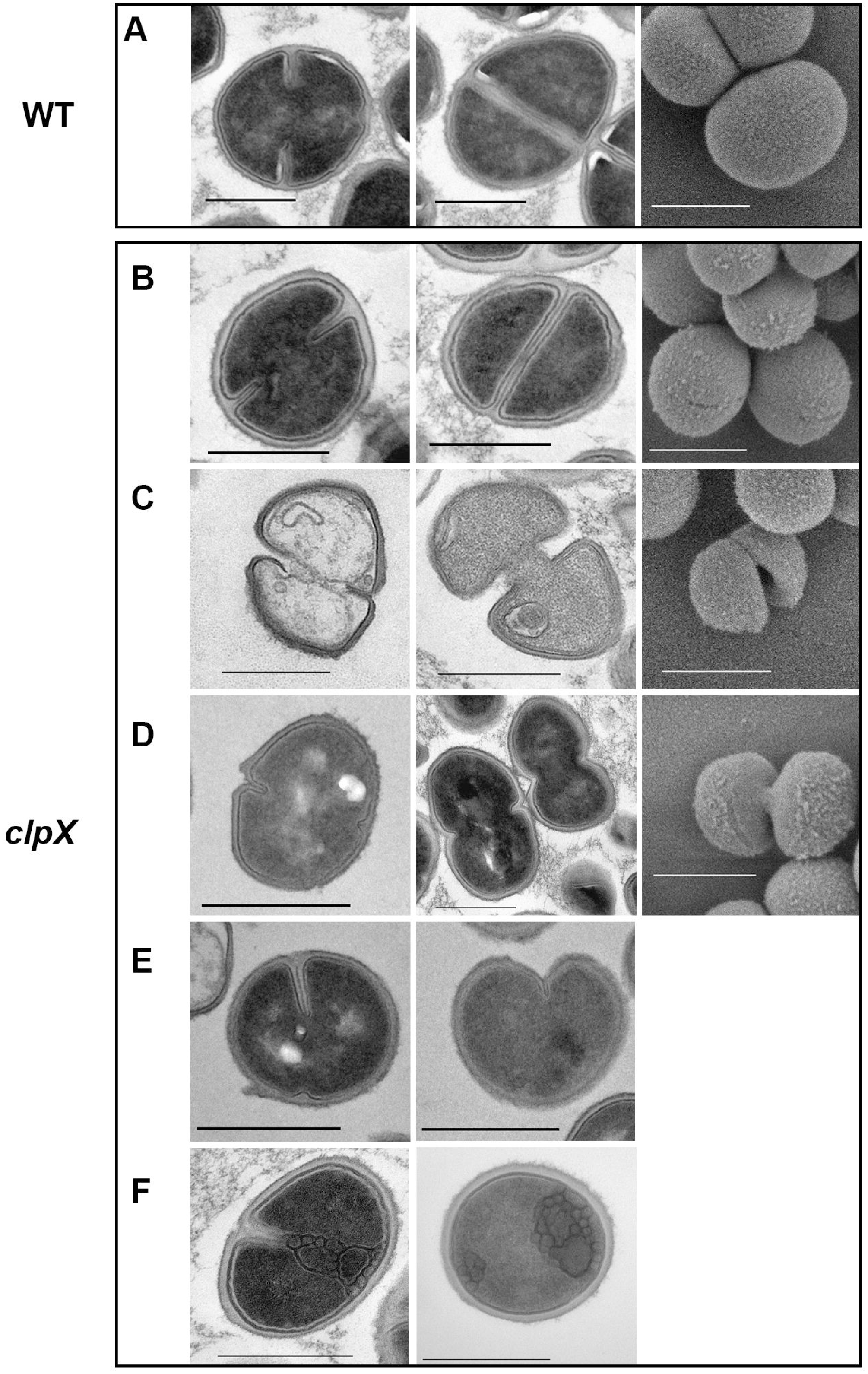
*S. aureus clpX* cells grown at 30°C display aberrant septum ingrowth and initiate daughter cell separation prior to septum closure. TEM (left panels) and SEM (right panel) images of wild-type (**a**) or *clpX* mutant cells (**b-f**) grown in TSB to mid-exponential phase at 30°C. Images show characteristic morphologies of wild-type or *clpX* cells at 30°C as determined from at least three biological replicates using different *S. aureus* strain backgrounds (SA564, 8325-4, and JE2 and the derived *clpX* mutants). The 1 μm scale-bar applies to TEM images in (**a**) and (**b**). Scale bars, 0.5 μm.

Some *clpX* cells also displayed septa extending asymmetrically towards the cell center, and in extreme cases extending inwards only from one side (+/- premature split; Fig 3d and 3e). This contrasts with wild-type *S. aureus* cells whose septa always extended symmetrically inwards from the edge of the cell wall (Fig 3a and S Fig 3). Finally, in some *clpX* cells, unordered membranous material reminiscent of mesosome-like structures [26] were observed at the site of septum ingrowth (Fig 3e and 3f). The latter phenotypes suggest that ClpX contributes to coordinating septum formation in *S. aureus*.

### The ClpX chaperone becomes critical for septum completion at 30°C

To quantify these phenotypes and to observe overall differences in progression of the cell cycle between wild-type and *clpX* cells, we performed Super-Resolution Structured Illumination Microscopy (SR-SIM) on cells stained with the membrane dye Nile red, and scored cells according to the stage of septum ingrowth as described by Monteiro et al. [24]; see Fig 4a for example images. To enumerate cells with incomplete septa that show signs of premature splitting, cells were additionally stained with fluorescently modified vancomycin (Van-FL), which labels the entire cell wall (cell periphery and septum), or with a green fluorescent derivative of wheat germ agglutinin WGA-488 that only labels the peripheral wall [24,27]. To estimate the number of lysed cells, DNA was stained with the blue dye Hoechst 3334. In this analysis, no significant differences in the distribution of cells in the different phases were observed for wild-type and *clpX* cells grown at 37°C (Fig 4a). At 30°C, however, significantly fewer *clpX* cells displayed a complete septum (4% as opposed to 15% of wild-type cells; P < 0.001). Moreover, while the fraction of cells that were in the process of building a septum (phase 2) was similar in wild-type and *clpX* cells at both temperatures, a more detailed analysis of the phase 2 cells revealed striking differences (Fig 4): consistent with the TEM analysis, a substantial number of *clpX* cells with incomplete septa showed signs of premature daughter cell splitting (20% of phase 2 cells), or had asymmetrical septum ingrowth (7% of phase 2 cells) when cells were grown at 30°C. None of these phenotypes were observed in wild-type cells at any temperature. While asymmetrical septum ingrowth was not observed in *clpX* cells grown at 37°C, premature splitting cells could be observed, however, at a lower frequency (Fig 4b). Furthermore, when subdividing phase 2 cells into two subclasses based on the extent of septum ingrowth, the proportion of *clpX* cells displaying early septum ingrowth (defined as cells with less than 15% septum ingrowth; see examples in Fig 4b) was significantly (P < 0.001) higher after growth at 30°C compared to 37°C, and when compared to the wild-type. For wild-type cells, an equal fraction of cells displayed early septum ingrowth at 30°C and 37°C. Finally, SR-SIM confirmed that the fraction of lysed *clpX* cells increased significantly when the temperature was decreased (2% at 37°C, and 16% at 30°C, P < 0.001). In comparison, the proportion of lysed wild-type cells was estimated to be below 2% at both temperatures.

**Fig 4.**
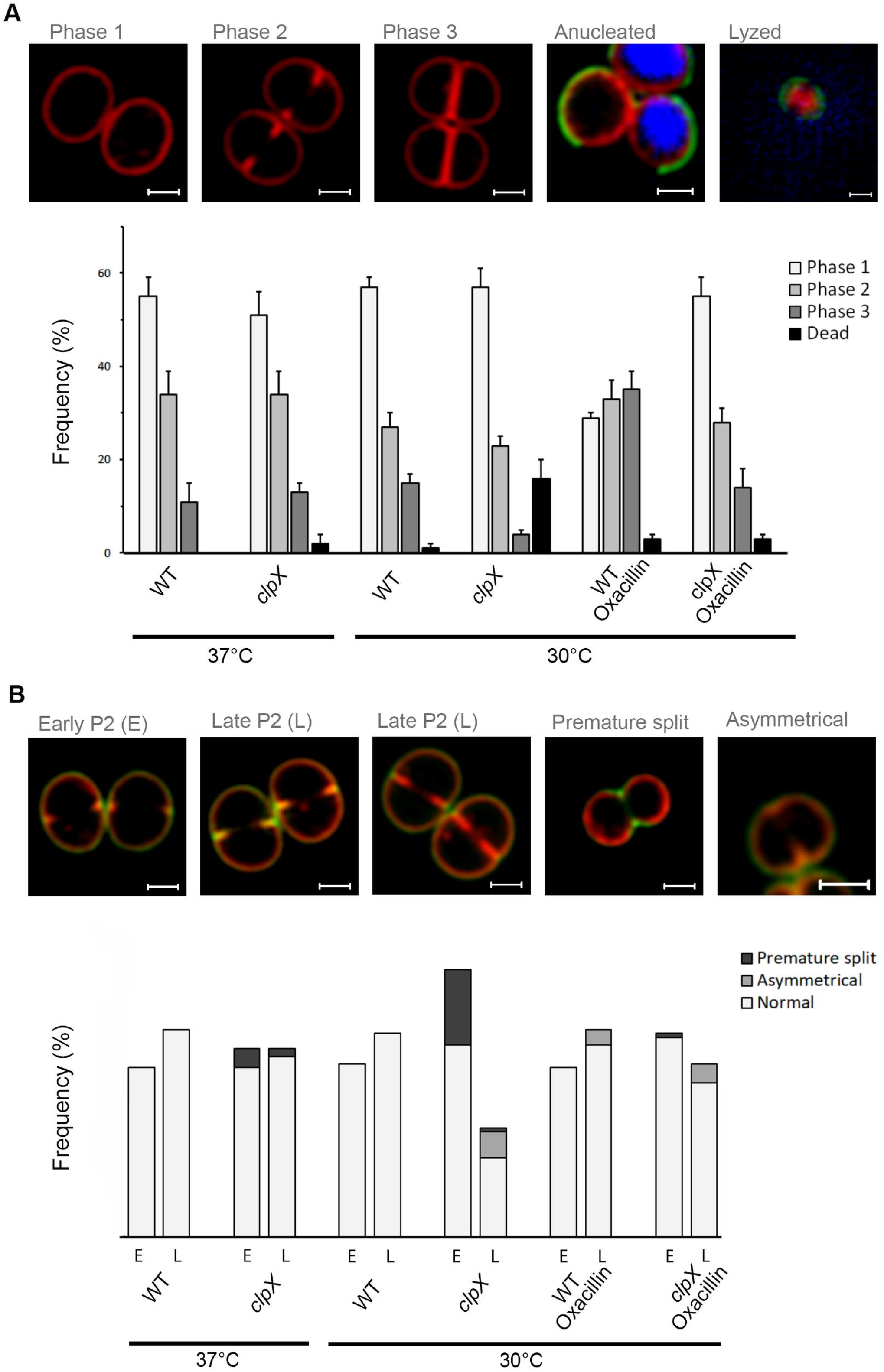
Oxacillin restores progression of the cell cycle in *S. aureus clpX* cells grown at 30°C. SA564 wild-type and *clpX* mutant cells were grown at 37°C or 30°C as indicated in the absence or presence of 0.05 μg ml^-1^ oxacillin (**a and b**); cells were then stained with membrane dye Nile Red (red) and cell wall dye Van-FL (green) before imaging by SR-SIM. To examine if ClpX alters progression of the growth cycle, 300 cells (from each of two biological replicates) were scored according to the stage of septum ingrowth: no septum (phase 1), incomplete septum (phase 2), or non-separated cells with complete septum (phase 3), according to the examples images shown in the (**a**) top panel. To enumerate the number of lysed cells, cells were additionally stained with the DNA dye Hoechst 3334. Scale bar, 0.5Lμm. (**b**) Phase 2 cells (200 phase 2 cells for each biological replicate) were additionally scored according to the state of septum ingrowth (cells with less than 15% septum ingrowth were scored as “early” (E), while cells with more than 15% septum ingrowth were scored as “late” (L)), and whether the ingrowth was asymmetrical, as shown in the example images in the top panel. The proportion of cells presenting premature split was estimated based on the Van-FL staining. Scale bar, 0.5Lμm.

In conclusion, the proportion of *clpX* mutant cells displaying a complete septum or late septum ingrowth was significantly reduced at 30°C, while the proportion of *clpX* cells displaying early septum ingrowth and aberrant septum was significantly increased at 30°C. Thus, the microscopy analyses suggest that ClpX chaperone activity becomes critical for the ability of *S. aureus* to complete the division septum as the temperature decreases.

### Oxacillin restores the cell cycle of *clpX* cells

To further examine how β-lactams improve growth of the *clpX* mutant, we performed SR-SIM analysis on oxacillin treated wild-type and *clpX* mutant cells grown at 30°C, as described above (Fig 4). Interestingly, sub-lethal concentrations of oxacillin significantly increased the fraction of phase 3 cells (closed septum): from 15 to 31% in the wild-type (P < 0.001), and from 4 to 14% in the *clpX* mutant (P < 0.001). Moreover, oxacillin significantly decreased the fraction of *clpX* cells (phase 2) that had initiated cell separation prior to septum completion from 20% to 2%, and in line with this observation, almost no lysed *clpX* mutant cells were observed (Fig 4b). Hence, oxacillin increases the fraction of cells with complete division septa in both the wild-type and the *clpX* backgrounds, and prevents premature splitting of *clpX* cells with incomplete division septa. In contrast, asymmetrical ingrowth of septa is still readily observed in oxacillin treated *clpX* mutant cells (Fig 4a and b).

These conclusions were supported when the oxacillin treated SA564 wild-type and *clpX* cells were analyzed by TEM (S4 Fig). TEM also showed that oxacillin prevented formation of mesosome-like structures in *clpX* cells (S4 Fig). However oxacillin treatment conferred a number of well described morphological changes that were shared by wild-type and *clpX* cells including thickening and misplacement of septa, a more fuzzy surface, and blurring of the electron-dense septal mid-zone (S4 Fig) [7,27,28]. Finally, many cells were present in pairs that have remained partly attached at the midline following septum completion (S4 Fig). Hence, both the SR-SIM and the TEM images support that oxacillin, even in concentrations well below the MIC value, prolongs phase 3 and delays splitting of the septum.

### Oxacillin antagonizes the septum progression defects conferred by inactivation of ClpX

To directly assess the impact of ClpX and oxacillin on progression of septal PG synthesis, we used an established metabolic labeling method with fluorescent D-amino acids (FDAAs) to visualize regions of new PG insertion [24,29,30]. PG synthesis was followed at 30°C and 37°C by sequentially labeling cells with FDAAs of different colors, thereby creating a virtual time-lapse image of PG synthesis [24,29,30]. Cells were first pulse-labeled for 10 min with green nitrobenzofurazan-amino-D-alanine (NADA), followed by a 10-min pulse with the blue hydroxycoumarin-amino-D-alanine (HADA). Labeled cells were imaged by SR-SIM, and progression of PG synthesis was scored in 300 randomly picked wild-type and *clpX* mutant cells grown in the absence or presence of oxacillin (Fig 5.,note that NADA is displayed in red). In the absence of oxacillin, PG synthesis proceeded from phase 1 (no septa, PG synthesis takes place in the lateral wall) to phase 2 (septal PG synthesis progresses inwards), and finally phase 3 (closed septum, PG synthesis occurs in both septum and the lateral wall) in > 95% of wild-type cells, as described in [24,25] (see Fig 5a). When the *clpX* mutant was grown at 37°C, PG synthesis followed the wild-type paradigm (S5 Fig). In contrast, when the *clpX* mutant was grown at 30°C, the septal PG synthesis progressed abnormally in a substantial fraction of phase 2 cells, as 22 ± 3 % of the *clpX* cells that had initiated septum formation in the first period of labeling (NADA) did not continue septum synthesis in the second period of labeling (HADA). Instead, the HADA signal co-localized with the NADA signal in the early septum ingrowth, and additionally, a peripheral HADA signal was visible (see examples in Fig 5a indicated by grey asterisks). Because other *clpX* cells displaying NADA labeling in an early septal ingrowth were indeed capable of septum progression and septum closure (green asterisks in Fig 5a), the septal PG synthesis rate does not seem to be generally reduced in the *clpX* mutant. Instead, the co-localization of the NADA and HADA in an early septum ingrowth may reflect stalling of septum synthesis in a subpopulation of *clpX* cells. Interestingly, in the presence of a sub-lethal concentration of oxacillin the fraction of *clpX* cells displaying co-localization of NADA and HADA at the early-septum ingrowth was reduced to 6 ± 2 % (Fig 5b).

**Fig 5.**
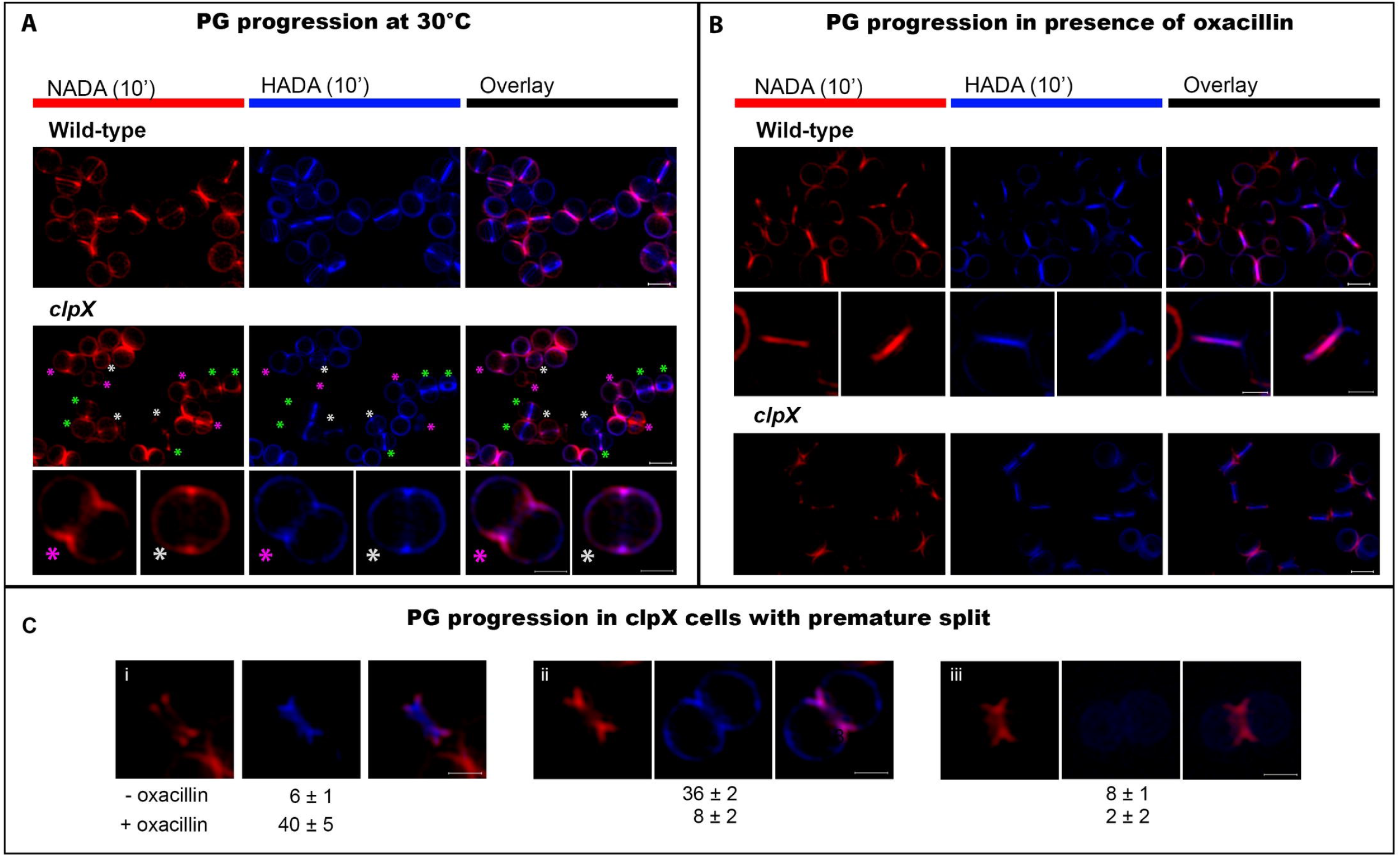
Aberrant progression of septal PG synthesis in *S. aureus* cells lacking ClpX is rescued by oxacillin. *S. aureus* wild-type (SA564) and *clpX* mutant cells were grown at 30°C and in the absence (**a** and **c**) or presence 0.05 μg ml^-1^ oxacillin (**b** and **c**), and PG synthesis was followed by sequentially labeling with NADA (green in primary data but recolored red to better distinguish it from the blue HADA signal) for 10 min, followed by washing and labeling with HADA for additional 10 min before SR-SIM imaging. (**a**) green asterisks mark cells displaying progression of septum synthesis = non-overlapping septal NADA and HADA signals; white asterisks mark cells with co-localization of NADA and HADA signals in an early septum ingrowth; pink asterisks mark cells displaying premature splitting. Lower panel shows enlarged examples of PG labeling in *clpX* cells displaying premature splitting and *clpX* cells where NADA and HADA signals overlap in an early septum ingrowth. (**b**) When *S. aureus* wild-type cells are grown in the presence of sub-lethal concentrations of oxacillin at 30°C, some cells display overlapping NADA and HADA septal signals, examples are shown in middle panel. (**c**) To examine progression of septal PG synthesis in *clpX* cells displaying premature split, 50 cells from each of three biological replicates (grown +/-oxacillin) that had initiated septum formation during incubation with NADA, and displayed premature splitting were randomly selected. PG synthesis was followed by assessing HADA incorporation. **(i-iii)** show examples and distribution of the three phenotypes observed. **(i)** show the number of cells where septum synthesis was continued; **(ii)** shows the number of cells where the HADA signal located to the peripheral wall; **(iii)** shows the number of cells where no HADA signal was detected. Numbers are given as the mean and SD of the three biological replicates. Scale bars, 0.5Lμm. *** P < 0.001; statistical analysis was performed using the chi square test for independence.

FDAAs only incorporate into newly synthesized PG and therefore premature splitting initiating from the peripheral wall cannot be detected with this approach [30]. However, splitting of newly synthesized, still incomplete, septum was observed (pink asterisks in Fig 5a), and while this phenotype was not observed in wild-type cells, this phenotype was displayed in about 20 ± 2 % of the *clpX* cells (phase 2 cells) grown in the absence of oxacillin. In the presence of oxacillin, only 8 ± 2 % of *clpX* cells showed splitting of newly synthesized still incomplete septa (see example in Fig 5b). In wild-type cells grown in the presence of oxacillin, NADA- and HADA signals more often co-localized in the entire septal plane (examples depicted in Fig 5b), supporting that wild-type cells grown with oxacillin spend longer time in phase 3. We conclude that at temperatures below the optimum, the ClpX chaperone activity becomes important for *S. aureus* septal PG synthesis to proceed beyond the point of septum initiation, and that oxacillin antagonizes the septum progression defects conferred by inactivation of ClpX.

### Oxacillin promotes septal PG synthesis in *clpX* cells with premature split

The results presented so far suggest that oxacillin improves growth of an *S. aureus clpX* mutant by stimulating septal PG synthesis and inhibiting premature splitting and lysis of daughter cells. To investigate septal PG synthesis in cells with premature splitting, we randomly picked 50 *clpX* cells grown at 30°C that had initiated septum formation during incubation with NADA, and that displayed the characteristic morphology of premature splitting, and assessed where HADA was incorporated in these cells. Interestingly, only very few *clpX* cells displaying premature septum split continued synthesizing septum (Fig 5c-i); instead HADA was incorporated at the cell periphery (Fig 5c-ii). In a few cells no HADA signal was detected at all (Fig 5c-iii). Similar results were obtained if the order of labeling was reversed (data not shown). Hence, septal PG synthesis seems to stop and instead become dispersed to the peripheral wall in *clpX* cells displaying splitting of a yet incomplete septum. Remarkably, in oxacillin treated cells, septum synthesis progressed normally in most cells with premature split (40 ± 1 of 50 cells, P < 0.001, Fig 5c). Taken together, this analysis demonstrates that oxacillin antagonizes the arrest of septum synthesis observed in *clpX* cells with premature septal split.

### FtsZ localization and dynamics are not affected in the *clpX* mutant

ClpX from diverse bacteria interacts directly with FtsZ suggesting that the ClpX chaperone has a conserved role in assisting assembly/disassembly of the FtsZ polymer [31-34]. We therefore reasoned that ClpX may regulate septum progression by interfering with FtsZ dynamics. To study localization and dynamics of FtsZ, a plasmid expressing eYFP-tagged derivative of FtsZ from an IPTG-inducible promoter [35] was introduced into *S. aureus* wild-type and *clpX* mutant cells, and time-lapse fluorescence microscopy was performed on cells growing on a semi-solid matrix at 30°C. In both wild-type and *clpX* mutant cells, Z-ring dynamics progressed predictably throughout the cell cycle (S2 Movie and Fig 6a): in newly divided cells, the Z ring has the same diameter as the cell until the ring starts to reduce in diameter and eventually closes (as described in [36]). Following closure, FtsZ undergoes a period of highly dynamic re-distribution, before the Z-ring cycle starts over again in newly divided cells. Hence, FtsZ dynamics appear not to be affected by lack of ClpX activity. Next, we imaged the relative localization of FtsZ and PG synthesis by sequentially labeling PG synthesis with FDAAs as described above, except that tetramethylrhodamine 3-amino–d-alanine (TADA, red signal) was used instead of NADA to avoid overlap with the yellow eYFP signal. In both wild-type and *clpX* cells, the eYFP signal localized ahead of septal PG synthesis in all phase 2 cells (Fig 6b and overview in S6 Fig). Specifically, FtsZ also localized ahead of the FDAA signal in *clpX* cells having HADA and TADA signal co-localizing in an early septum in growth (Fig 6b). Strikingly, the FtsZ signal maintained its septal localization in *clpX* cells with premature split and arrest of septal PG-synthesis (see example in Fig 6b). As also shown above, PG incorporation in such cells takes place in the peripheral wall. Hence, our data supports the idea that FtsZ dynamics is not impeded in cells lacking ClpX.

**Fig 6.**
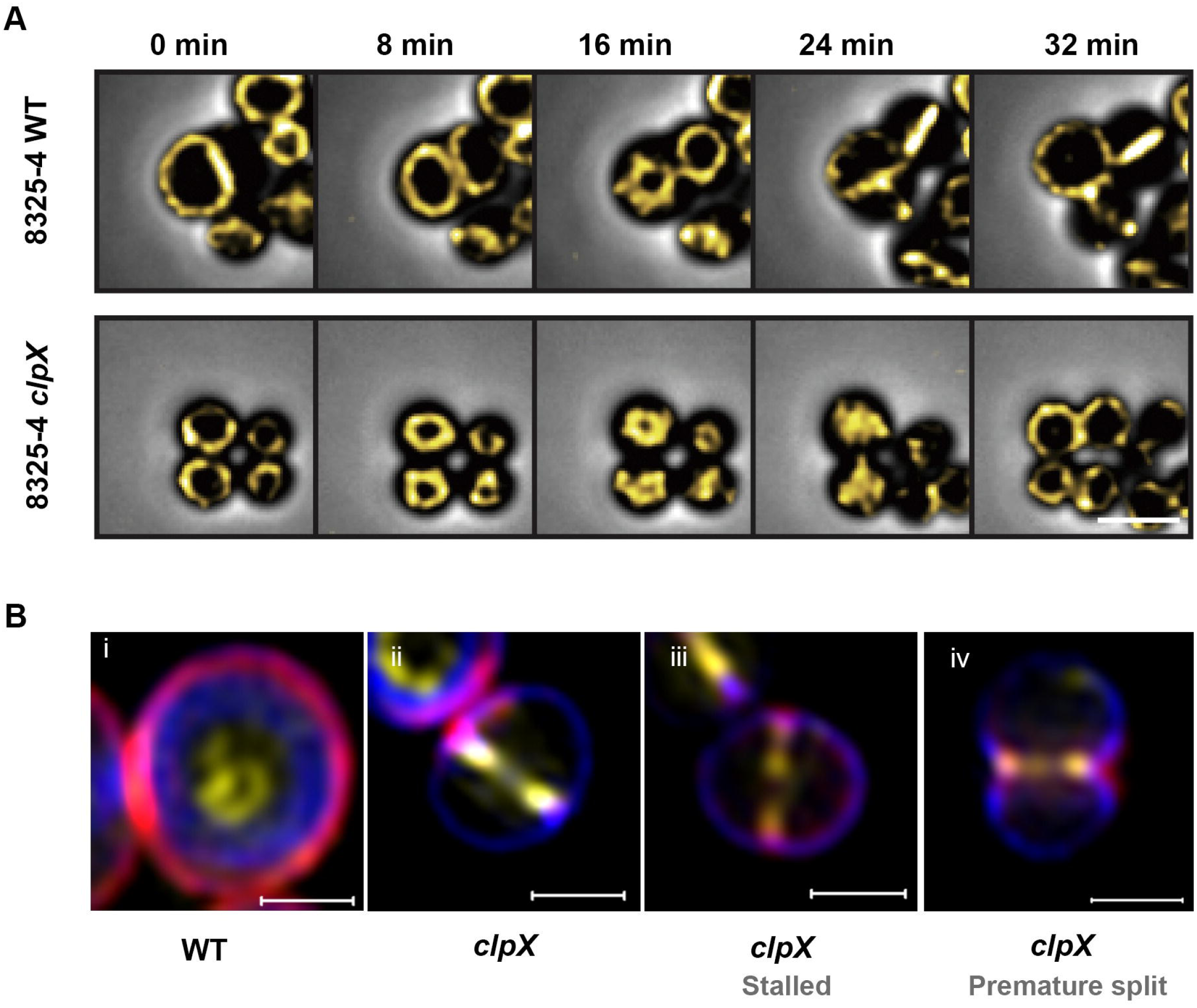
FtsZ localization and dynamics appear similar in *S. aureus* wild-type and *clpX* mutant cells. FtsZ localization and dynamics were analyzed in *S. aureus* (8325-4) wild-type and *clpX* mutant expressing an eYFP-tagged derivative of FtsZ expressed from an IPTG-inducible promoter. (**a**) Still images from time-lapse fluorescence microscopy showing FtsZ dynamics in *S. aureus* wild type and *clpX* cells growing on a semi-solid matrix in the presence of 100 uM IPTG at 30°C (S2 Movie). The fluorescent signal is overlaid with the phase contrast. Scale bar 1 μm.(**b**) Localization of FtsZ relative to PG synthesis was analyzed by sequentially FDAA labeling *S. aureus* wild-type and *clpX* cells growing in TSB with + 50 uM IPTG at 30°C: TADA (red) for 10 minutes followed by washing and labeling with HADA (blue) for additional 10 min prior to SR-SIM imaging. Overview images can be found in S6 Fig. Examples of cells that started septum synthesis in the first period of labeling displaying a septal FtsZ-eYFP signal ahead of the site of active PG synthesis (**i**) wild-type cell (**ii**) *clpX* cell (**iii**) *clpX* cell with co-localization of TADA and HADA signals; and (**iv**) *clpX* cell displaying premature split. Scale bars 0.5 μm.

### Inhibitors of WTA biosynthesis rescue growth of the *clpX* mutant

Finally, we asked if the ability to rescue growth of an *S. aureus clpX* mutant is specific for the β-lactam class of antibiotics (S7 Fig). The compounds assessed were either antibiotics with completely different targets, or compounds inhibiting various steps in the cell envelope synthesis pathway. Interestingly, only tunicamycin and tarocin A1, two well characterized inhibitors of WTA biosynthesis that work synergistically with β-lactams to kill *S. aureus* [14,16], stimulated growth of the *clpX* mutant (Fig 7 and S7 Fig). Strikingly, tunicamycin was almost as efficient as oxacillin in stimulating growth of the *clpX* (Fig 7c). In contrast, other late stage inhibitors of PG synthesis, such as vancomycin, did not stimulate growth of the *clpX* mutant, even though this antibiotic, similarly to β-lactams, prevents PG cross-linking (S7 Fig). Lysostaphin, which breaks already formed cross-bridges, also had no stimulatory effect on growth of the *clpX* mutant (S7 Fig). These findings indicate that reduced cross-linking *per se* does not alleviate the growth defect of the *clpX* mutant and that the growth defect of *S. aureus clpX* mutants is specifically rescued by tunicamycin, tarocin A1, and β-lactam antibiotics underscoring a functional link between the PBP activity and WTA biosynthesis.

**Fig 7.**
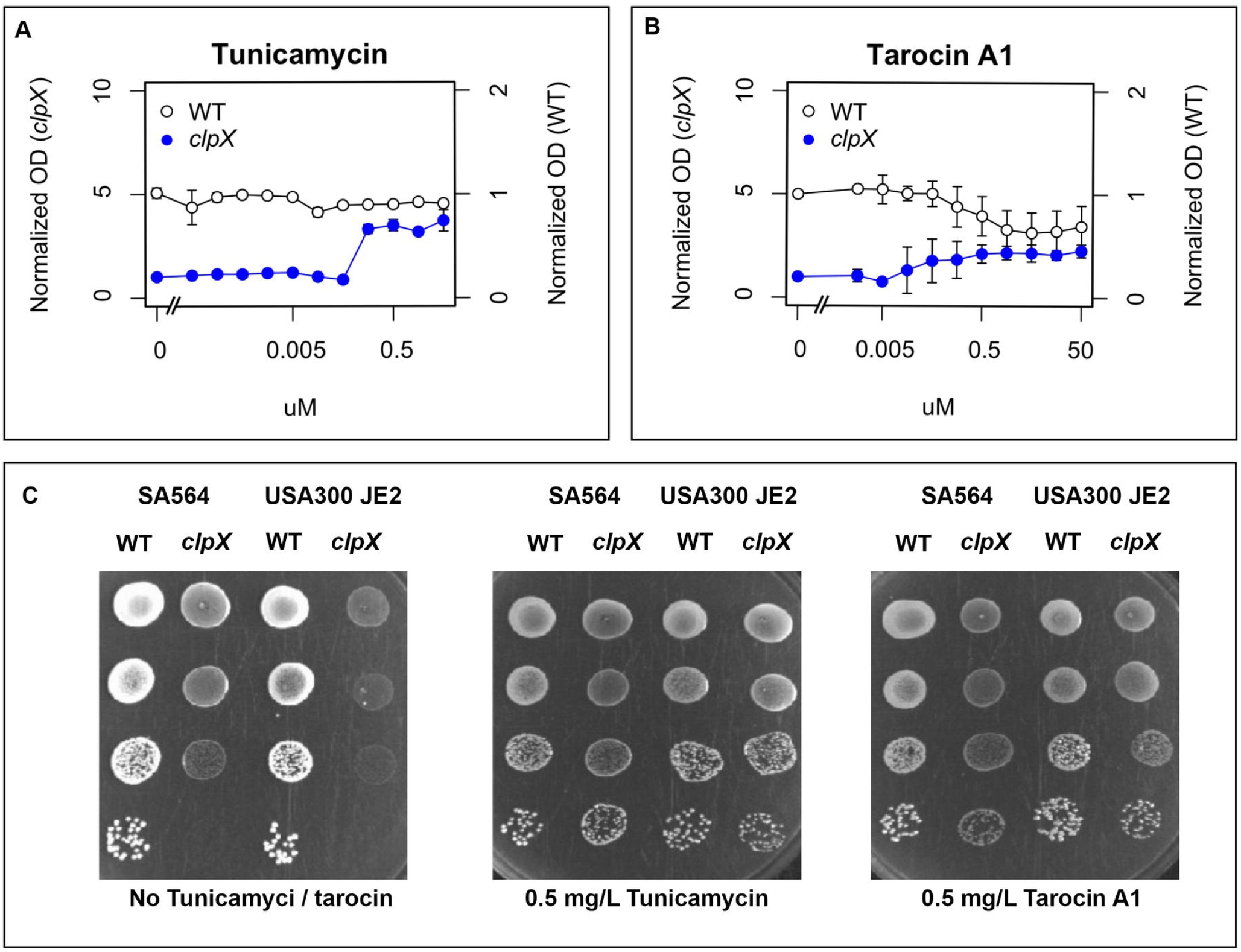
Growth of *S. aureus clpX* mutants is stimulated by inhibitors of WTA synthesis. (**a** and **b**): *S. aureus* SA564 wild type and *clpX* strains were grown overnight at 37°C, diluted 1:200 and grown at 37°C until mid-exponential phase. These cultures were then diluted into TSB containing increasing concentrations of the indicated compounds in a 96-well format, and the plates were incubated for 24 h at 30°C. The values represent means of OD values, normalized to the OD values obtained without compound. Error bars indicate standard deviations. Note that different scales were used on the two axes due to the difference in growth between the WT and *clpX* mutant: values for the *clpX* mutant are indicated on the left vertical axis, and values for the WT are indicated on the right vertical axis to allow easy comparison of growth between the two strains. (**c** and **d**) The *S. aureus* wild-type strains, SA564 (MSSA) and USA300 JE2 (MRSA) and the corresponding *clpX* mutants were grown exponentially in TSB at 37°C. At OD_600_ = 0.5, cultures were diluted 10^1^, 10^2^, 10^3^ and 10^4^-fold, and 10 μl of each dilution was spotted on TSA plates in the presence or absence of tunicamycin/tarocin A1 (as indicated) and incubated at 30°C for 24 h.

## Discussion

Assembly of the bacterial cell division machinery is a highly coordinated process with proteins recruited to the division site in a specific order and depending on the timely interaction between a large number of proteins [37]. Here, we show that the widely conserved ClpX chaperone plays an important role in staphylococcal cell division at 30°C but not at 37°C. In wild-type *S. aureus* cells, splitting of daughter cells is not initiated prior to septum closure. In contrast, a substantial fraction of *clpX* cells displaying incomplete septa had initiated splitting of daughter cells indicating that the system responsible for coordinating autolytic splitting with septum completion has become dysregulated. In *clpX* cells displaying the premature splitting phenotype, septal PG synthesis was not continued, and instead became dispersed to the peripheral wall demonstrating that *clpX* cells with premature split are unable to finalize the septum. The detrimental character of this defect likely prevents cells from undergoing further divisions, explaining why a large proportion of *clpX* cells are non-dividing and end up lysing. In support hereof, TEM pictures show that most *clpX* ghost cells were in the process of splitting despite having an incomplete septum. This is likely due to turgor pressure forces breaking the tip of the ingrowing septum where the cell wall is thin and mechanically weak [38]. Hence, we assume that premature splitting is the underlying cause for the high rate of spontaneous lysis observed among *clpX* cells.

Importantly, cells devoid of ClpX contain elevated levels of the two major autolysins associated with separation of *S. aureus* daughter cells, Sle1 and Atl [20,21,39,40]. Therefore, premature splitting of *clpX* cells could simply be a consequence of excess autolysins. However, whilst the elevated levels of SleI and Atl may contribute to the premature splitting and spontaneous lysis of *clpX* cells, we believe that additional factors are in play. This is based on the findings that i) premature splitting and lysis of *clpX* cells is more frequent at 30°C than at 37°C, whereas autolysin levels are elevated at both temperatures [20,21,39,40]; and, ii) the *clpX* phenotypes described here are not shared by a *clpP* mutant, although *clpP* mutant cells also contain elevated Sle1 levels [20,21,23,41]. Taken together these findings indicate that the high levels of autolysins are more detrimental to *clpX* cells growing at 30°C, suggesting that ClpX in addition to controlling the levels of autolysins also affects their activation via a temperature dependent pathway. As stalling of early septum synthesis in *clpX* cells was observed only at the lower temperature, we speculate that stalling of septum synthesis contributes to premature activation of autolysins in the *clpX* mutant as depicted in the working model shown in Fig 8. In this model, *S. aureus* depends on ClpX chaperone activity for transforming an early stage divisome complex into a late stage divisome complex at 30°C but not at 37°C. At both temperatures, the high levels of autolysins in the *clpX* mutant will make the mutant more prone to initiate daughter cell separation before septum completion. However, stalling of the divisome by an unknown mechanism exacerbates the risk of premature split at 30°C. In support hereof, premature split was also observed in *clpX* cells grown at 37°C, however, at this temperature septal progression seems to proceed fast enough to enable completion of the septum, as outlined in Fig 8. Notably, FtsZ localization and dynamics were not affected in the absence of ClpX, suggesting that ClpX affects septum formation and autolytic activation downstream of Z-ring formation. Intriguingly, *S. aureus* cytokinesis was recently proposed to occur in two-steps: an initial FtsZ dependent slow step that may drive the initial membrane invagination, and a second faster step driven by PG synthesis and recruitment of late division proteins such as MurJ [36]. Hence, we speculate that ClpX promotes septum formation at 30°C by assisting recruitment of MurJ or other central components of the late divisome complex.

**Fig 8.**
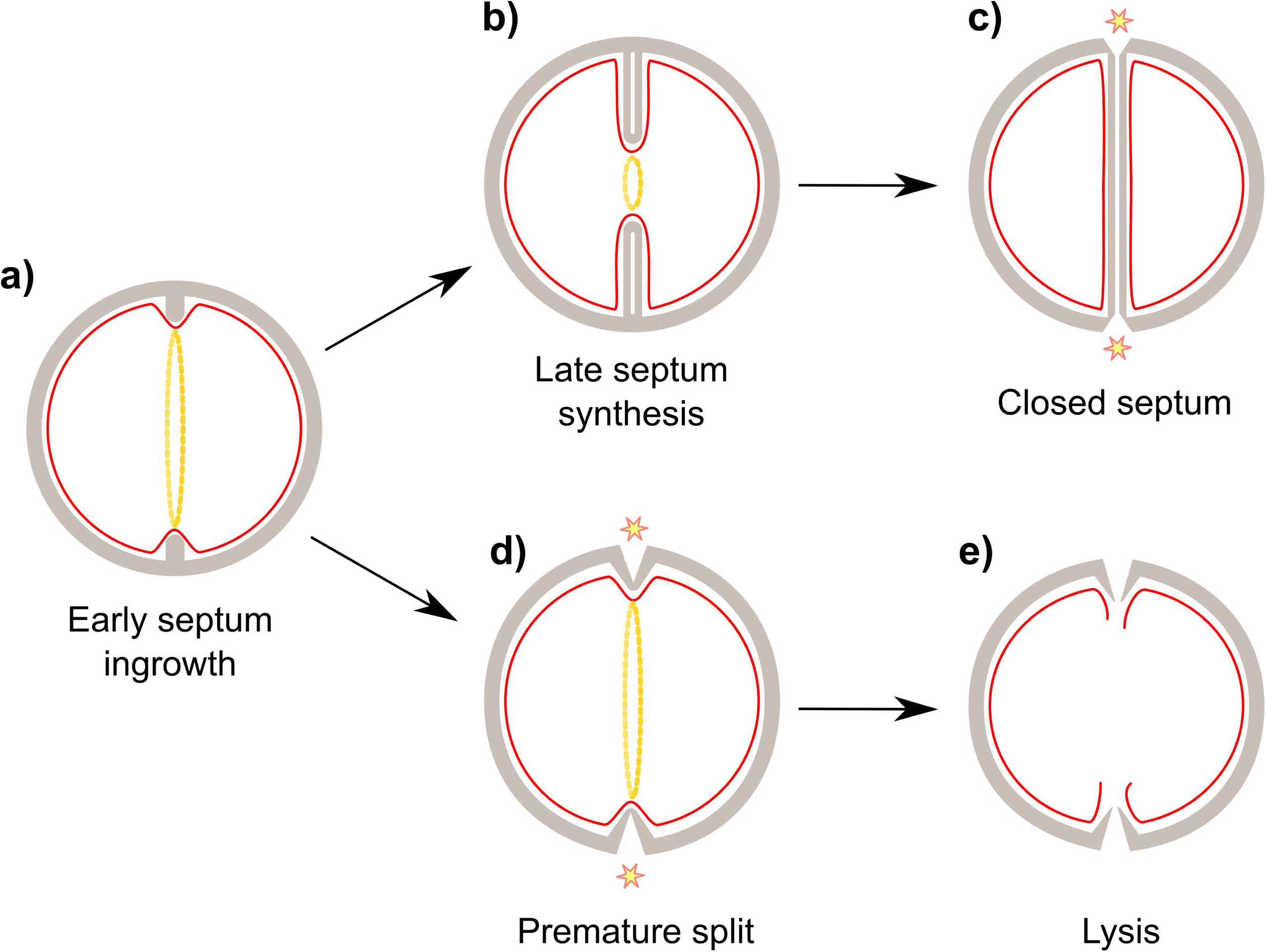
Model of temperature dependent *S. aureus* cell division. Progression from an early septal ingrowth (**a**) to a late septal ingrowth (**b**) is dependent on protein-rearrangements in the divisome. At 37°C these rearrangements occur spontaneously, however at 30°C these rearrangements need assistance from the ClpX chaperone. (**c**) Upon septum closure, autolytic activity is activated. (**d**) In *S. aureus* cells lacking ClpX activity, septum synthesis will stall in an early septal ingrowth leading to activation of autolytic splitting from the peripheral wall and eventually (**e**) cell lysis unless autolytic activation is inhibited by β-lactams.

Because mis-coordination in activation of autolytic enzymes may have fatal consequences for the cell, regulatory checkpoints that coordinate the autolytic system with septum completion likely exist, however, little is known about these mechanisms. Remarkably, the growth and lysis defect of the *clpX* mutant was alleviated by sub-lethal concentrations of β-lactam antibiotics. This intriguing finding is to our knowledge the first example of β-lactam antibiotics increasing the growth-rate and preventing spontaneous lysis of a bacterial mutant. The presented data show that oxacillin simultaneously rescues septum synthesis, and prevents premature splitting, mesosome formation, and spontaneous lysis of the *clpX* mutant, lending further support to a linkage between these phenotypes. The ability of sub-lethal concentrations of β-lactam antibiotics to suppress spontaneous lysis of *clpX* mutant cells was surprising, as the lethal activity of β-lactam antibiotics is believed to stem from the loss of wall integrity accompanied by cell lysis [5,9,42]. Here, we observed that oxacillin treatment of both *S. aureus* wild-type and *clpX* mutant cells increased the fraction of cells displaying a complete division septum, supporting previous findings that β-lactams delay autolytic splitting of daughter cells [7,28]. Moreover, the sequential PG staining experiments showed that late septal FDAA signals often overlap in wild-type cells grown in the presence of oxacillin, indicating that β-lactams also prolong PG synthesis in the completed septum (which may be a consequence of delayed autolytic cell splitting). Consistent with these findings, β-lactams treated *S. aureus* cells display characteristic thickened septum in TEM images [7,28]. One possible scenario to explain how β-lactams rescue *clpX* cells is therefore that oxacillin prevents spontaneous lysis of the *clpX* mutants by suppressing activation of autolytic enzymes and by stimulating late septal PG synthesis (Fig 8). Vice versa, the earlier onset of autolytic activity mediated by inactivation of ClpX, may counteract the delay in autolytic splitting of daughter cells observed in oxacillin treated wild-type cells, thereby explaining the reduced cell generation time of *clpX* cells compared to wild-type cells in the presence of oxacillin.

In support of a central role of autolysins in the *clpX* phenotypes, we previously showed that the fast-growing suppressor mutants arising when *clpX* cells are grown at 30°C have loss-of-function mutations in the *ltaS* gene encoding the LtaS synthetase that is essential for synthesis of LTA, another key regulator of autolytic activity [20]. Similarly, WTA seems to impede splitting of the septal cross wall and control Atl localization [14,43]. Strikingly, two inhibitors of WTA synthesis, tunicamycin and tarocin A1, were the only other compounds that similarly to β-lactams rescued growth of the *clpX* mutant. In our working model, we therefore propose that inhibition of WTA stimulates growth of the *clpX* mutant by impeding premature split of *clpX* cells (Fig 8).

In conclusion, we have shown that *S. aureus* cell division is temperature sensitive, and that at 30°C, the ClpX chaperone serves an important function in coordinating initiation of daughter cell separation with septum completion. When ClpX is absent, cell division frequently has a fatal outcome because septal PG synthesis stalls and cell separation is initiated prior to completion of the septum. Interestingly, these defects were prevented by binding of β-lactam antibiotics to the PBP transpeptidase activity domain, indicating that this final stage in PG biosynthesis plays a role in coordinating septum synthesis and activation of autolytic splitting of daughter cells. Consistent with this hypothesis, the transpeptidase site of PBP1 was proposed to take part in a checkpoint-type mechanism functioning at the end of each round of cell division to ensure that autolytic splitting of daughter cells can only take place upon successful completion of septum synthesis [44,45]. Our work therefore supports the idea that in this clinically important bacterium, the effect of β-lactam antibiotics is tightly linked to coordination of cell division.

## Materials and Methods

### Bacterial strains and growth conditions

Strains used in this study are listed in S2 Table. *S. aureus* strains were grown in tryptic soy broth media (TSB; Oxoid) under vigorous agitation at 200 rpm at 37°C. In most experiments, 20 ml of medium was inoculated in 200-ml flasks to allow efficient aeration of the medium. For solid medium, 1.5% agar was added to make TSA plates. Erythromycin (7.5 μg ml^-1^) was added as required. Upon receipt of the low-passage isolate SA564, the strain was cultured once and stored frozen at -80°C. In all experiments, we used bacterial strains freshly streaked from the frozen stocks on TSA plates with antibiotics added as required and incubated overnight at 37°C. The growth was followed by measuring the optical densities at 600 nm. The starting OD was always below 0.05. When inoculating *S. aureus clpX* deletion strains, care was taken to avoid visibly larger colonies containing potential suppressor mutants [20]. To reduce the selection pressure for suppressor mutants in broth cultures, strains were upon inoculation first grown at 37°C for four generations (OD_600_ ~0.1-0.2) before shifting to 30°C. *S. aureus* JE2-derived strains were obtained from the Network of Antimicrobial Resistance in Staphylococcus aureus (NARSA) program (supported under NIAID/NIH contract HHSN272200700055C.

### Growth calculations

Growth of *S. aureus* strains was assessed by measuring optical density (OD_600_) of cultures grown in 96-well microplates (for end-point OD) or in a Bioscreen C instrument (for growth rates). Overnight cultures were grown in TSB at 37°C. For end-point ODs, overnight cultures were diluted 1:200 in TSB and grown to exponential phase (OD_600_ 0.1) and then diluted 1:10,000 in 200 μl TSB in 96-well format, and incubated 24 h at 30°C with shaking. For growth in the Bioscreen C instrument overnight cultures were diluted in 300 μl TSB to an OD_600_ of approx. 0.001, and OD_600_ was measured every 5 min with 20 seconds of shaking before each measurement. All exponential growth rates were determined by growing the relevant strains in a Bioscreen C instrument as described above. The growth rates were automatically calculated as described before [20]. In short, OD_600_ values were log-transformed and linear regressions were determined for each data point in the OD_600_ interval from 0.02 to 0.12 based on a window containing 15 data points. The exponential growth rate was identified as the maximal slope of the linear regressions. The standard error of the mean was calculated using values from different biological replicates.

### Disc diffusion assays

*S. aureus* strains were inoculated on TSA plates and incubated at 37°C overnight. The next day, a bacterial suspension was adjusted to a 0.5 McFarland (Sensititre^®^ nephelometer and the Sensititre^®^ McFarland Standard) and streaked on MHA. The plates were allowed to dry prior to the addition of 1 μg oxacillin discs (Oxoid) and incubated at 37°C for 48 hours.

### Time-lapse microscopy

*S. aureus* wild-type (SA564 or 8325-4/pCQ11ftsZ::eYFP) and *clpX* mutant (SA564ΔclpX or 8325-4ΔclpX/pCQ11ftsZ::eYFP) were grown overnight in TSB medium at 37°C and cultures were diluted 100 times in fresh TSB medium and grown until an OD_600_ of 0.1. Cells were washed once in fresh TSB medium and spotted onto a TSB-polyacrylamide (10%) slide incubated with TSB medium supplemented when appropriate with 0.008 μg ml^-1^ oxacillin or with 100 μM IPTG. Acrylamide pads were placed inside a Gene frame (Thermo Fisher Scientific) and sealed with a cover glass as described before [46].

Phase contrast images acquisition was performed using a DV Elite microscope (GE healthcare) with a sCMOS (PCO) camera with a 100x oil-immersion objective. Images were acquired with 200 ms exposure time every 2 or 4 minutes for at least 4 h at 30 °C using Softworx (Applied Precision) software. Images were analyzed using Fiji (http://fiji.sc). Each experiment was performed at least in triplicate. 69 micro colonies were imaged for SA564; 63 micro colonies for SA564 + oxacillin; 43 micro colonies for SA564 *clpX* and 36 micro colonies for SA564 *clpX +* oxacillin. Cell tracking was performed using the TrackMate plugin (https://imagej.net/TrackMate) in Fiji, and the output was converted to Newick format for lineage tree plotting and calculation of cell generation times using a custom R script (available at http://github.com/ktbaek/beta-lactams-clpx).

Time-lapse images of FtsZ-eYFP were acquired using a Leica DMi8 microscope with a sCMOS DFC9000 (Leica) camera with a 100x oil-immersion objective and a Spectra X (Lumencor) illumination module. Fluorescent images were acquired every 4 min with 400 ms exposure using a YFP filter cube (Chroma, excitation 492-514 nm, dichroic 520 nm, emission 520-550 nm). Images were processed using LAS X (Leica) and signal was deconvolved using Huygens (SVI) software.

### Electron microscopy and image analysis

Overnight cultures grown at 37°C were diluted 1:200 into 40 ml of fresh TSB and grown at 30°C or 37°C to an OD_600_ of 0.5. Bacteria (SA564, 8325-4, and JE2 and the *clpX* mutant derived here from) from a 10-ml culture aliquot were collected by centrifugation at 8,000 x g, and the cell pellets were suspended in fixation solution (2.5% glutaraldehyde in 0.1 M cacodylate buffer [pH 7.4]) and incubated overnight at 4°C. The fixed cells were further treated with 2% osmium tetroxide, followed by 0.25% uranyl acetate for contrast enhancement. The pellets were dehydrated in increasing concentrations of ethanol, followed by pure propylene oxide, and then embedded in Epon resin. Thin sections for electron microscopy were stained with lead citrate and observed in a Philips CM100 BioTWIN transmission electron microscope fitted with an Olympus Veleta camera with a resolution of 2,048 by 2,048 pixels. For quantitative analysis, the images were acquired in an unbiased fashion by using the multiple image alignment function in the ITEM software (Olympus). Sample processing and microscopy were performed at the Core Facility for Integrated Microscopy (CFIM), Faculty of Health and Medical Sciences, University of Copenhagen.

### Scanning electron microscopy (SEM)

Exponentially growing *S. aureus* SA564 were collected by centrifugation, fixed in 2% glutaraldehyde in 0.05 M sodium phosphate buffer, pH 7.4 and sedimented on coverslips for 1 week at 4?°C. The cells were washed three times with sodium cacodylate buffer and progressively dehydrated by immersion in a graded series of ethanol (50–100%). Cells were subsequently mounted on stubs using colloidal silver as an adhesive, and sputter coated with gold (Leica Coater ACE 200) before imaging with a FEI Quanta 3D scanning electron microscope operated at an accelerating voltage of 2 kV. Sample preparation and SEM imaging was performed at CFIM.

### SR-SIM analysis

SR-SIM was performed with an Elyra PS.1 microscope (Zeiss) using a Plan-Apochromat 63x/1.4 oil DIC M27 objective and a Pco.edge 5.5 camera. Images were acquired with five grid rotation and reconstructed using ZEN software (black edition, 2012, version 8.1.0.484) based on a structured illumination algorithm, using synthetic, channel specific optical transfer functions and noise filter settings ranging from -6 to -8. Laser specifications can be seen in S3 Table. SR-SIM was performed at CFIM.

### Analysis of the cell cycle

To address progression of the cell cycle, exponential cultures of *S. aureus* were incubated for 5 min at room temperature with the membrane dye Nile Red, the cell wall dye WGA-488 or Van-Fl and the DNA dye Hoechst 3334 (S4 Table). Samples were placed on an agarose pad and visualized by SR-SIM as described above. 300 cells were scored according to the stage of septum ingrowth: no septum (phase 1), incomplete septum (phase 2), or non-separated cells with complete septum (phase 3). Dead cells were scored based on Hoechst staining: lysed cells, as cells where DNA had leaked out of the cell and anucleated cells as cells devoid of Hoechst staining. The analysis was performed on two biological replicates.

Additionally, 200 cells were scored according to the state of septum ingrowth (cells with less than 15% septum ingrowth were scored as “early”, while cells with more than 15% septum ingrowth were scored as “late”) and whether the ingrowth was asymmetrical or showed signs of premature splitting. The latter was based on staining with Van-FL. This analysis was performed on two biological replicates.

### Analysis of progression of PG synthesis

To evaluate localization of PG synthesis, exponential cultures of *S. aureus* (SA564 or 8325-4) were pulse labeled with FDAAs; cells were initially incubated 10 minutes with NADA, washed in PBS and resuspended in TSB. The cells were then incubated 10 minutes with HADA, washed with PBS, placed on an agarose pad and visualized by SR-SIM. This experiment was conducted in three biological replicates including a staining in reverse order and one using the red TADA as a replacement for NADA. Analysis on the progression of PG synthesis was performed on 300 cells for each biological replicate with similar results.

To investigate the progression of septal PG synthesis in *clpX* mutant cells displaying premature split, HADA incorporation was assessed in 50 cells (in each of three biological replicate) that had initiated septum formation during the initial labeling and displayed the characteristic morphology of premature splitting were selected randomly. PG synthesis was followed by assessing.

### FtsZ localization in SR-SIM

In order to assess FtsZ relative to the active PG synthesis, *S. aureus* (8325-4) wild-type and *clpX* mutant transformed with pCQ11 expressing an eYFP-tagged derivative of FtsZ from an IPTG-inducible promoter were analyzed using sequentially labeling with FDAAs as described above (incubation with TADA for 10 minutes followed by HADA for 10 minutes). Cells were grown at 30°C in the presence of 50 μM IPTG (at higher IPTG concentrations cell division defects were observed in the wild-type strain).

### Statistical analysis

Statistical analysis was done using R statistical software. Student’s t-test was used to assess significant differences in growth in the absence or presence of a tested antibiotic. The Chi-squared test of independence was used to determine if there was a significant relationship between the proportion of cells assign to each of the three phases or relevant phenotypes under the tested condition (number of cells in the relevant phase or phenotype/the total number of cells). A value P < 0.05 was considered significant.

## Acknowledgments

We would like to thank Professor Simon Foster (University of Sheffield) for the generous gift of FDAA’s and the FtsZ-eYFP fusion plasmid. We would like to thank Ewa Kuninska (University of Copenhagen) for excellent technical assistance and the staff at the Core Facility for Integrated Microscopy (University of Copenhagen) for their enthusiastic assistance in doing SEM, TEM and SR-SIM.

## Supporting information

**S1 Fig. β-lactams stimulate growth of a *S. aureus clpX* deletion strains, but not of wild type or *clpP* deletion strains.**

Mean growth rates (h^-1^) for SA564, JE2, and Newman strains and corresponding *clpX* and *clpP* deletion mutants grown in the presence of increasing concentrations of β-lactams at 30°C. Numbers indicate mean doubling time in minutes. The standard error of the mean (error bars) was calculated using values from three biological replicates. Asterisks indicate significantly improved growth (P < 0.05). The P values were obtained by comparing the growth rates at each antibiotic concentration to the growth rate without antibiotics and were calculated using Student’s t-test

**S2 Fig. Tracking of single *S. aureus* cells in micro-colonies**

**a**. Micro-colony cell lineage trees. Each cell in a representative micro-colony of each strain and condition was tracked for the first 8h of the time-lapse experiment shown in Fig 2a, using the TrackMate plugin in Fiji. The diagram shows the resulting cell lineage trees. Red dots indicate cell lysis. **b**. Single-cell generation time. Each point represents a newly divided daughter cell with generation time (time until next division) plotted on the vertical axis, and elapsed time at cell birth plotted on the horizontal axis.

**S3 Fig. Morphological changes in *S. aureus clpX* cells grown at 30°C.**

TEM images of *S. aureus* SA564 wild type cells (upper panel) and SA564 *clpX* cells (lower panel) harvested in exponential phase at 30°C. Note the many lyzed cells in TEM images of the *clpX* mutant. Scale bar, 5.0 μm.

**S4 Fig. Oxacillin delays separation of daughter cells.**

TEM images of SA564 wild-type (panel a and c) or *clpX* cells (panel b and d) cells grown in TSB to mid-exponential phase at 30°C in the absence (panel a and b) or presence of 0.05 mg/L oxacillin (panel c and d). The scale bar corresponds to 1.0 μm. The images show several features of β-lactam treated wild-type and *clpX* cells such as a weak or missing midline arrow (arrow A), a fuzzy cell wall appearance (arrow B), and cells failing to separate after division (arrow C). The asymmetrical septum ingrowth can still be observed in oxacillin treated the *clpX* mutant cells (arrow D).

**S5 Fig. PG synthesis in *S. aureus clpX* cells grown at 37°C follows the wild-type paradigm.**

SA564*clpX* cells were grown at 37°C in the absence of oxacillin and PG synthesis was followed by sequentially labeling with TADA (red) for 10 min, followed by washing and labeling with HADA (blue) for additional 10 min before cells were imaged using SR-SIM. Scale bar 1.0 μm. The red and blue signals do not overlap, illustrating that septal peptidoglycan synthesis is progressing predictably inwards and PG synthesis follows the wild-type paradigm for cells in phase 1, 2 and 3. Images shown are representative of three biological replicates.

**S6 Fig. FtsZ localization relative to PG synthesis in wild-type and clpX mutant**

FtsZ localization was analyzed in *S. aureus* wild-type and *clpX* cells expressing an eYFP-tagged derivative of FtsZ expressed from an IPTG-inducible promoter. Localization of FtsZ relative to PG synthesis was analyzed by sequentially labeling *S. aureus* wild type and *clpX* cells growing in TSB supplemented with 50 uM IPTG at 30°C with TADA (red) for 10 minutes followed by washing and labeling with HADA (blue) for additional 10 min prior to SR-SIM imaging. Images shown are representative of cells from three biological replicates. Scale bars, 1 μm (overview), 0.5 μm (single cells).

**S7 Fig. Effect of different antibiotics on growth of the wild type and *clpX* cells**

*S. aureus* SA564 wild type and *clpX* strains were grown overnight at 37°C, diluted 1:200 and grown at 37°C until mid-exponential phase. These cultures were then diluted into TSB containing increasing concentrations of the indicated compounds in a 96-well format, and the plates were incubated for 24 h at 30°C. The values represent means of OD values, normalized to the OD values obtained without compound. Error bars indicate standard deviations. Note that different scales were used on the two axes due to the difference in growth between the WT and *clpX* mutant: values for the *clpX* mutant are indicated on the left vertical axis, and values for the WT are indicated on the right vertical axis to allow easy comparison of growth between the two strains.

## References

1. DeLeo FR, Otto M, Kreiswirth BN, Chambers HF. Community-associated meticillin-resistant *Staphylococcus aureus*. Lancet. 2010;375: 1557–1568.

2. Elander RP. Industrial production of beta-lactam antibiotics. Appl Microbiol Biotechnol. 2003;61: 385–392.

3. Wise EM, Park JT. Penicillin: its basic site of action as an inhibitor of a peptide cross-linking reaction in cell wall mucopeptide synthesis. Proc Natl Acad Sci USA. 1965;54: 75–81.

4. Tipper DJ, Strominger JL. Mechanism of action of penicillins: a proposal based on their structural similarity to acyl-D-alanyl-D-alanine. Proc Natl Acad Sci USA. 1965;54: 1133–1141.

5. Tomasz A. The mechanism of the irreversible antimicrobial effects of penicillins: how the beta-lactam antibiotics kill and lyse bacteria. Annu Rev Microbiol. 1979;33: 113–137.

6. McDowell TD, Reed KE. Mechanism of penicillin killing in the absence of bacterial lysis. Antimicrob Agents Chemother. 1989;33: 1680–1685.

7. Giesbrecht P, Kersten T, Maidhof H, Wecke J. Staphylococcal cell wall: morphogenesis and fatal variations in the presence of penicillin. Microbiol Mol Biol Rev. 1998;62: 1371–1414.

8. Bayles KW. The bactericidal action of penicillin: new clues to an unsolved mystery. Trends Microbiol. 2000;8: 274–278.

9. Cho H, Uehara T, Bernhardt TG. Beta-lactam antibiotics induce a lethal malfunctioning of the bacterial cell wall synthesis machinery. Cell. 2014;159: 1300–1311.

10. Pinho MG, Kjos M, Veening J-W. How to get (a)round: mechanisms controlling growth and division of coccoid bacteria. Nat Rev Microbiol. 2013;11: 601–614.

11. Reed P, Atilano ML, Alves R, Hoiczyk E, Sher X, Reichmann NT, et al. *Staphylococcus aureus* survives with a minimal peptidoglycan synthesis machine but sacrifices virulence and antibiotic resistance. PLoS Pathog. 2015;11: e1004891.

12. Peacock SJ, Paterson GK. Mechanisms of methicillin resistance in *Staphylococcus aureus*. Annu Rev Biochem. 2015;84: 577–601.

13. Rolo J, Worning P, Nielsen JB, Sobral R, Bowden R, Bouchami O, et al. Evidence for the evolutionary steps leading to *mecA*-mediated β-lactam resistance in staphylococci. PLoS Genetics. 2017;13: e1006674.

14. Campbell J, Singh AK, Santa Maria JP Jr, Kim Y, Brown S, Swoboda JG, et al. Synthetic lethal compound combinations reveal a fundamental connection between wall teichoic acid and peptidoglycan biosyntheses in *Staphylococcus aureus*. ACS Chem Biol. 2011;6: 106–116.

15. Farha MA, Leung A, Sewell EW, D’Elia MA, Allison SE, Ejim L, et al. Inhibition of WTA synthesis blocks the cooperative action of PBPs and sensitizes MRSA to β-lactams. ACS Chem Biol. 2012;8: 226–233.

16. Lee SH, Wang H, Labroli M, Koseoglu S, Zuck P, Mayhood T, et al. TarO-specific inhibitors of wall teichoic acid biosynthesis restore β-lactam efficacy against methicillin-resistant staphylococci. Sci Transl Med. 2016;8: 329ra32.

17. Drawz SM, Bonomo RA, Three decades of β-lactamase inhibitors. Clin Microbiol Rev. 2010;23: 60–201.

18. Olivares AO, Baker TA, Sauer RT. Mechanistic insights into bacterial AAA+ proteases and protein-remodelling machines. Nat Rev Microbiol. 2016;14: 33–44.

19. Frees D, Qazi S, Hill PJ, Ingmer H. Alternative roles of ClpX and ClpP in *Staphylococcus aureus* stress tolerance and virulence. Mol Microbiol. 2003;48:1565–1578.

20. Bæk K, Bowman L, Søgaard M, Kaever V, Siljamäki P, Savijoki K, et al. The cell wall polymer lipoteichoic acid becomes non-essential in *Staphylococcus aureus* cells lacking the ClpX chaperone. MBio. 2016;7: e01228–16.

21. Stahlhut SG, Alqarzaee AA, Jensen C, Fisker NS, Pereira AR, Pinho MG, et al. The ClpXP protease is dispensable for degradation of unfolded proteins in *Staphylococcus aureus*. Sci Rep. 2017;7: 11739.

22. Percy MG, Gründling A. Lipoteichoic acid synthesis and function in gram-positive bacteria. Annu Rev Microbiol. 2014;68: 81–100.

23. Bæk KT, Gründling A, Mogensen RG, Thøgersen L, Petersen A, Paulander W, et al. β-Lactam resistance in methicillin-resistant *Staphylococcus aureus* USA300 is increased by inactivation of the ClpXP protease. Antimicrob Agents Chemother. 2014;58: 4593–4603.

24. Monteiro JM, Fernandes PB, Vaz F, Pereira AR, Tavares AC, Ferreira MT, et al. Cell shape dynamics during the staphylococcal cell cycle. Nat Commun. 2015;6: 8055.

25. Zhou X, Halladin DK, Rojas ER, Koslover EF, Lee TK, Huang KC et al. Bacterial division. Mechanical crack propagation drives millisecond daughter cell separation in *Staphylococcus aureus*. Science. 2015;348: 574–8.

26. Kahl BC, Belling G, Reichelt R, Herrmann M, Proctor RA, Peters G. Thymidine-dependent small-colony variants of *Staphylococcus aureus* exhibit gross morphological and ultrastructural changes consistent with impaired cell separation. J Clin Microbiol. 2003;41:410–3.

27. Paul TR, Venter A, Blaszczak LC, Parr TR, Labischinski H, Beveridge TJ. Localization of penicillin-binding proteins to the splitting system of *Staphylococcus aureus* septa by using a mercury-penicillin V derivative. J Bacteriol. 1995;177: 3631–3640.

28. Lorian V. Some effect of subinbilitory concentrations of penicillin on the structure and division of staphylococci. Antimicrob Agents Chemother. 1975;7: 864–867.

29. Kuru E, Hughes HV, Brown PJ, Hall E, Tekkam S, Cava F, et al. In situ probing of newly synthesized peptidoglycan in live bacteria with fluorescent D-amino acids. Angew Chem Int Ed. 2012;51: 12519–12523.

30. Lund VA, Wacnik K, Turner RD, Cotterell BE, Walther CG, Fenn SJ, et al. Molecular coordination of *Staphylococcus aureus* cell division. Elife. 2018;7: e32057.

31. Weart RB, Nakano S, Lane BE, Zuber P, Levin PA. The ClpX chaperone modulates assembly of the tubulin-like protein FtsZ. Mol Microbiol. 2005;57: 238–249.

32. Dziedzic R, Kiran M, Plocinski P, Ziolkiewicz M, Brzostek A, Moomey M, et al. Mycobacterium tuberculosis ClpX Interacts with FtsZ and Interferes with FtsZ Assembly. PLoS ONE. 2010;5: e11058.

33. Sugimoto S, Yamanaka K, Nishikori S, Miyagi A, Ando T, Ogura T. AAA+ chaperone ClpX regulates dynamics of prokaryotic cytoskeletal protein FtsZ. J Biol Chem. 2010;285: 6648–6657.

34. Haeusser DP, Lee AH, Weart RB, Levin PA. ClpX inhibits FtsZ assembly in a manner that does not require its ATP hydrolysis-dependent chaperone activity. J Bacteriol. 2009;191: 1986–1991.

35. Liew AT, Theis T, Jensen SO, Garcia-Lara J, Foster SJ, Firth N, et al. A simple plasmid-based system that allows rapid generation of tightly controlled gene expression in *Staphylococcus aureus*. Microbiology. 2011;157: 666–76.

36. Monteiro JM, Pereira AR, Reichmann NT, Saraiva BM, Fernandes PB, Veiga H, et al. Peptidoglycan synthesis drives an FtsZ-treadmilling-independent step of cytokinesis. Nature. 2018;554:528–532.

37. Haeusser DP, Margolin W. Splitsville: structural and functional insights into the dynamic bacterial Z ring. Nat Rev Microbiol. 2016;14: 305–19.

38. Matias VR, Beveridge TJ. Native cell wall organization shown by cryo-electron microscopy confirms the existence of a periplasmic space in *Staphylococcus aureus*. J Bacteriol. 2006;188: 1011–1021.

39. Yamada S, Sugai M, Komatsuzawa H, Nakashima S, Oshida T, Matsumoto A, et al. An autolysin ring associated with cell separation of *Staphylococcus aureus*. J Bacteriol. 1996;178: 1565–1571.

40. Kajimura J, Fujiwara T, Yamada S, Suzawa Y, Nishida T, Oyamada Y, et al. Identification and molecular characterization of an N-acetylmuramyl-l-alanine amidase Sle1 involved in cell separation of *Staphylococcus aureus*. Mol Microbiol. 2005;58: 1087–1101.

41. Feng J, Michalik S, Varming AN, Andersen JH, Albrecht D, Jelsbak L, et al. Trapping and proteomic identification of cellular substrates of the ClpP protease in *Staphylococcus aureus*. J Proteome Res. 2013;12: 547–58.

42. Chung HS, Yao Z, Goehring NW, Kishony R, Beckwith J, Kahne D. Rapid beta-lactam-induced lysis requires successful assembly of the cell division machinery. Proc Natl Acad Sci USA. 2009;106: 21872–21877.

43. Schlag M, Biswas R, Krismer B, Kohler T, Zoll S, Yu W, et al. Role of staphylococcal wall teichoic acid in targeting the major autolysin Atl. Mol Microbiol. 2010;75: 864–73.

44. Pereira SFF, Henriques AO, Pinho MG, Tomasz A. Role of PBP1 in cell division of *Staphylococcus aureus*. J Bacteriol. 2007;189: 3525–3531.

45. Pereira SFF, Henriques AO, Pinho MG, Lencastre H de, Tomasz A. Evidence for a dual role of PBP1 in the cell division and cell separation of *Staphylococcus aureus*. Mol Microbiol. 2009;72: 895–904.

46. de Jong IG, Beilharz K, Kuipers OP, Veening JW. J. Live Cell Imaging of *Bacillus subtilis* and *Streptococcus pneumoniae* using Automated Time-lapse Microscopy. Vis Exp. 2011;53.

